# Dynamic functional connectivity in the right temporoparietal junction captures variations in autistic trait expression

**DOI:** 10.1101/2023.06.05.543654

**Authors:** Laura Bravo Balsa, Ahmad Abu-Akel, Carmel Mevorach

## Abstract

**Background:** Autistic individuals can experience difficulties with attention reorienting and Theory of Mind (ToM), which are closely associated with anterior and posterior subdivisions of the right temporoparietal junction. While the link between these processes remains unclear it is likely subserved by a dynamic crosstalk between these two subdivisions. We therefore examined the dynamic functional connectivity (dFC) between the anterior and posterior TPJ, as a biological marker of attention and ToM, to test its contribution to the manifestation of autistic trait expression in Autism Spectrum Disorder (ASD).

**Methods:** Two studies were conducted, exploratory (14 ASD, 15 TD) and replication (29 ASD, 29 TD), using resting-state fMRI data and the Social Responsiveness Scale (SRS) from the ABIDE repository. Dynamic Independent Component Analysis was performed in both datasets using the CONN toolbox. An additional sliding-window analysis was performed in the replication study to explore different connectivity states (from highly negatively to highly positively correlated).

**Results:** dynamic FC was reduced in ASD compared to TD adults in both the exploratory and replication datasets and was associated with increased SRS scores (especially in ASD). Additional regression analyses revealed that for ASD, decreased SRS autistic expression was predicted by engagement of highly negatively correlated states, while engagement of highly positively correlated states predicted increased expression.

**Conclusions:** These findings provided consistent evidence that the difficulties observed in ASD are associated with altered patterns of dFC between brain regions subserving attention reorienting and ToM processes.

## Introduction

Autism spectrum disorder (ASD) is a neurodevelopmental disorder characterised by difficulties in social interaction, communication, and repetitive, restricted behaviours (1). Within the wide spectrum of autistic traits, atypicalities in Theory of Mind (ToM)/mentalising and attention reorienting, including the possible link between them, have been reported by an extensive body of behavioural and neuroimaging research (2–5). ToM refers to the ability to understand and infer the mental states - beliefs, intentions, emotions - of oneself and others, and to use this information to predict behaviour (6–8). Previous studies have consistently reported difficulties in the behavioural performance of ASD groups compared to typically developing (TD) controls during ToM tasks (such as false beliefs tasks) (9,10). These behavioural effects are also mirrored in the decreased activation and reduced functional connectivity (FC) in ASD of the neural network underpinning ToM (11–14), which chiefly consists of the medial prefrontal cortex (mPFC), the temporal pole, the posterior cingulate cortex (PCC), and the temporo-parietal junction (TPJ).

Attention atypicality in orienting, disengaging and reorienting of attention when presented with novel relevant stimuli (i.e., “attention shifting” or “attention reorienting”) have also been consistently demonstrated in ASD (15,16). For instance, using a peripheral cuing paradigm (17), Keehn and colleagues (2) showed that autistic participants were significantly slower than TD participants in orienting their attention to a validly cued target. Likewise, several studies have found that ASD participants shifted their attention towards both social and non-social stimuli less frequently than TD participants (18,19). Here too, neuroimaging studies in ASD have shown atypical activation in regions associated with attention orienting and reorienting, mainly within the ventral frontoparietal attention network. Reduced activation of the inferior frontal gyrus and the supramarginal gyrus in response to the appearance of social cues were recorded in ASD (20), as well as both under- and over-reactivity in prefrontal cortex and the inferior parietal lobule for passive or active novelty detection, respectively (21,22).

The ability to shift attention to new relevant stimuli both in social and non-social contexts could be a pre-requisite for infants’ development of ToM (23), premised on the notion that shifting attention between different spatially distinct external stimuli is similar to the requirement to shift attention between your own and another person’s point of view (24, 25). Evidence for the association of early manifestation of attention disengagement difficulties (between 6 and 12 months) and later diagnosis of ASD at 24 months (26) has increased the interest in the link between ToM and attention difficulties in ASD (15,27,28).

The involvement of the right TPJ (rTPJ) in both reorienting attention and ToM processes have been consistently documented, (27–32). For example, brain stimulation over the rTPJ has been shown to impair both attention shifting performance (using a visual cueing paradigm) and ToM performance (increased error rates in a false belief task) (33). A closer inspection of the rTPJ using diffusion-weighted imaging tractography–based parcellation (31) found that the right anterior TPJ (raTPJ) is connected with the ventral prefrontal cortex and anterior insula (parts of the attention network), while the right posterior TPJ (rpTPJ) shows stronger connectivity with the posterior cingulate cortex (PCC), temporal pole and medial prefrontal cortex (parts of the ToM network). Corresponding with this structural parcellation, Krall et al. (34) reported functional evidence showing higher activation of the raTPJ during attention reorienting tasks and higher activation of the rpTPJ during ToM tasks.

Subsequent resting-state FC studies have revealed antagonistic relationship between the raTPJ and rpTPJ (28,30,35,36). For example, using coactivation-based parcellation, Bzdok et al. (30) found the attention network to be both positively correlated with the raTPJ and anti-correlated with the rpTPJ; whereas the social cognition network showed positive correlation with the rpTPJ and negative correlation with the raTPJ. Accordingly, the authors proposed that the posterior and anterior rTPJ belong to two antagonistic brain networks (30). It is evident that two distinct and possibly antagonistic brain networks, with a focus in the TPJ, support attention orienting and ToM. It is, therefore, possible that an important proxy for understanding manifestation of autistic expression, which is associated with atypicalities in both of these processes, can be found in the functional connectivity between the raTPJ (attention) and rpTPJ (ToM) parcellations of the right TPJ region.

Previous FC studies in ASD are inconsistent and tend to split between a hypo (37,38) and hyper (39,40) connectivity pattern in the social cognition network of ASD participants compared to neurotypical individuals. These inconsistencies may point to the limitations of static FC studies especially in the context of neural networks underlying complex cognitive processes, such as attention and ToM (41–43). In contrast to the traditional static FC, *dynamic* FC tracks the temporal variations in FC resulting in different brain state configurations (connectivity patterns) among neural networks at different time points (41,42,44). Thus, for instance, at the beginning of a scan, a pair of regions of interest (ROIs) could show a high positive correlation, whereas later in the scan they could be negatively correlated. Existing dynamic FC research demonstrates its utility in several domains, including for predicting behaviour (45), classifying bipolar disorder (46), and detecting abnormal connectivity patterns associated with dysfunctional thoughts in major depression disorder (47) and with clinical symptoms in schizophrenia (48).

The limited extant research using dynamic FC studies in ASD has successfully highlighted group differences particularly in the average time spent in each state of connectivity (i.e., mean dwell time, MDT) between ASD and TD participants (49). In a recent paper (40), both group differences in overall dynamic FC variability in some regions (e.g., in the saliency and default mode networks) and differences in specific connectivity states (specifically the time spent in and the likelihood of transitioning into a hyper-connected state) were documented, and were also associated with autistic traits on the individual level (40). Thus, dynamic FC investigations are important in ASD in both highlighting global differences in the overall dynamic nature of connectivity (i.e., whether connections remain constant or change during rest) and in the specific connectivity states that occur during rest, including the time a person ‘spends’ in a particular state or the likelihood they will transition into that state.

In the current study, we focus on the dynamic resting state FC between the anterior and posterior subdivisions of the rTPJ as a window into the relationship between attention reorienting and social cognition in autistic compared to TD individuals (28). For this purpose, dynamic independent component analysis (dyn-ICA) was used on two resting-state fMRI datasets from the Autism Brain Imaging Data Exchange (ABIDE) database (http://fcon_1000.projects.nitrc.org/indi/abide/) (50,51), with the first serving as an exploratory and the second as a replication. dyn-ICA was used to assess general variability in the functional connectivity (i.e., a measure of the degree to which functional connectivity changes throughout the resting-state scan). In addition, we assessed specific connectivity states (from highly negatively correlated to highly positively correlated) in the larger replication dataset using a sliding-window approach (47). The latter enables us to identify the underlying source of the global functional connectivity variability difference. Previous research argued that integration processes would be enhanced by higher FC variability (52,53). Since we understand attention orienting to be intrinsically linked to ToM (23), we expect the two networks to be supported by a degree of functional connectivity variability, and therefore atypicalities in attention and ToM (as in ASD) should be associated with reduced variability. In our analysis this should be evidenced in decreased dynamic resting-state FC between the raTPJ and rpTPJ in the ASD group. In addition, as previous research pointed to the antagonistic relation between the networks —although this is in static connectivity— we expected ASD participants to show reduced likelihood of entering into a high negative correlation state of the raTPJ and rpTPJ. Finally, both the reduction in variability and the reduced likelihood of negative states are expected to predict increased manifestation of autistic traits on the individual level, which may support the dimensional notion of ASD (40,48,49,54).

## Methods

### Participants

Original fMRI and phenotypic data were obtained from the open-access ABIDE repository (http://fcon_1000.projects.nitrc.org/indi/abide/) (50,51). A first exploratory study was conducted using the University of Leuven Sample 1 dataset (subsequently referred to as Exploratory), which includes the resting-state fMRI data of 14 high-functioning autistic males (21.7 ± 4.0 years) and 15 typically developing (TD) matched controls (23.3 ± 2.9 years). A second study was conducted using the Barrow Neurological Institute dataset (subsequently referred to as Replication) to replicate and expand the first findings in a larger cohort. The Replication dataset includes the resting-state fMRI data of 29 high-functioning autistic males (37.4 ± 16.1 years) and 29 TD matched controls (39.6 ± 15.1 years).

### Clinical assessment

Across both datasets, autistic traits expression was assessed with the self-report Social Responsiveness Scale (SRS) (55), which significantly discriminated between groups. The SRS measures social difficulties related to ASD and quantifies their levels of expression. The scale includes a total score and five subscales that assess social awareness, social cognition, social communication, social motivation, and autistic mannerisms. Higher SRS scores indicate higher expression of autistic traits. Demographics and clinical characteristics are available in Table 1 in the Results section.

**Table 1.** Demographic characteristics and autistic trait expression (both datasets)

|  | Exploratory |  |  | Replication |  |  |
| --- | --- | --- | --- | --- | --- | --- |
|  | ASD<br>(*n* = 14)<br>Mean ± SD | TD<br>(*n* = 15)<br>Mean ± SD | *p*-value | ASD<br>(*n* = 29)<br>Mean ± SD | TD<br>(*n* = 29)<br>Mean ± SD | *p*-value |
| Age (years) | 21.7 ± 4.0 | 23.3 ± 2.9 | 0.223 | 37.4 ± 16.1 | 39.6 ± 15.1 | 0.604 |
| Sex | All males | All males |  | All males | All males |  |
| FIQ | 107.9 ± 13.9 | 114.8 ± 12.8 | 0.160 | 107.8 ± 13.7 | 112.4 ± 12.1 | 0.175 |
| SRS – Total (self-report) | 76.9 ± 24.2 | 43.6 ± 21.7 | <0.001 | 106.3 ± 30.4 | 27.9 ± 17.9 | <0.001 |
| *SRS subscales* |  |  |  |  |  |  |
| Social awareness | 9.2 ± 3.1 | 6.8 ± 2.8 | 0.03 | 11.4 ± 4.5 | 5.2 ± 2.8 | <0.001 |
| Social cognition | 14.5 ± 4.8 | 7.2 ± 4.8 | <0.001 | 19.7 ± 6.3 | 4.7 ± 4.3 | <0.001 |
| Social communication | 25 ± 6.9 | 14.4 ± 8.1 | <0.001 | 35.8 ± 10.9 | 7.6 ± 6.6 | <0.001 |
| Social motivation | 15 ± 4.8 | 7.9 ± 4.8 | <0.001 | 18.9 ± 6.3 | 6.4 ± 4.8 | <0.001 |
| Autistic mannerisms | 13.1 ± 5.1 | 7.1 ± 4.5 | 0.002 | 20.3 ± 6.8 | 4.1 ± 3.6 | <0.001 |

Abbreviations: FIQ, Full scale IQ; SRS, Social Responsiveness Scale. Higher scores in SRS represent higher autistic trait expression.

### Data acquisition

Data acquisition parameters for the Exploratory and Replication datasets are available in Supplementary Material and were obtained from http://fcon_1000.projects.nitrc.org/indi/abide:

ABIDE I University of Leuven Sample 1 and ABIDE II Barrow Neurological Institute repositories, respectively.

### Data preprocessing and Denoising

Functional data was analysed in the CONN toolbox 19.c (https://www.nitrc.org/projects/conn/) (56), an open-source software based on Statistical Parametric Mapping (SPM12, https://www.fil.ion.ucl.ac.uk/spm/) and MATLAB (MathWorks, Natick, MA). Data preprocessing and denoising was performed using batch scripts (available online at https://github.com/laurabravo97/Preprocessing-Denoising-CONN-toolbox-) following the CONN default pipelines (57) with minimal adjustments. Please see Supplementary Material for details on preprocessing and denoising steps.

### Dynamic Independent Component Analysis (Dyn-ICA)

Two regions of interest (ROI) masks for the raTPJ and the rpTPJ (see Figure 1A) were obtained from Mars et al. (31) and are available at http://www.rbmars.dds.nl/CBPatlases.htm. Fslmaths (FMRIBS’s Software Library, https://fsl.fmrib.ox.ac.uk, version 6.0.1) was used to remove the overlap between them. ROI-to-ROI dynamic resting-state FC analysis was performed using the dyn-ICA approach available in the CONN toolbox. Dyn-ICA is a data-driven approach that returns spatial maps containing connectivity patterns that can be interpreted as building blocks of dynamic FC (57). In this case, because two seeds are introduced in the analysis, we obtain one component/circuit timeseries that explains most of the variance across the 2×2 dynamic FC matrix.

**Figure 1.**
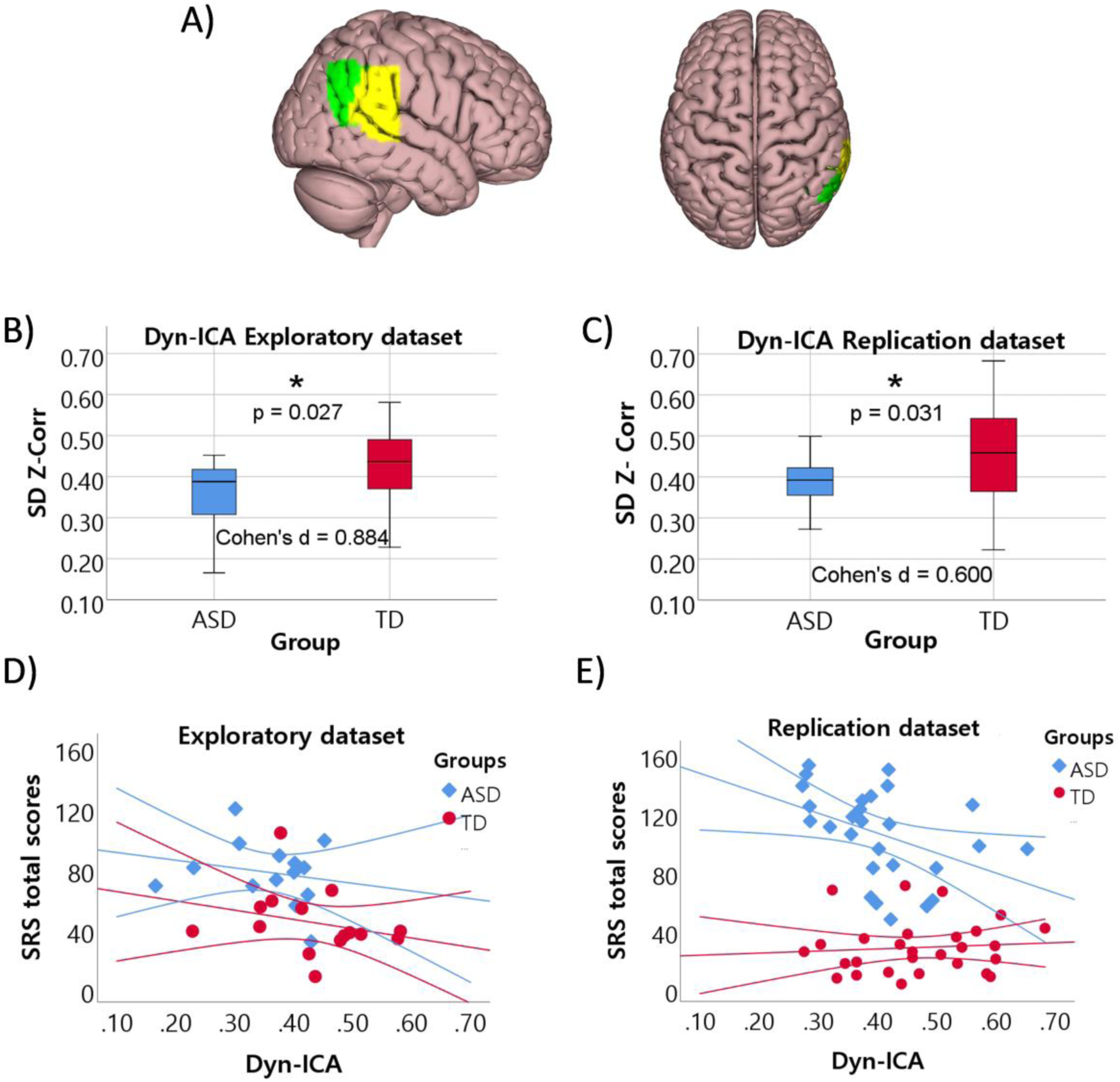
Dynamic resting-state functional connectivity (FC) between the anterior and posterior subdivisions of the right temporoparietal junction (raTPJ and rpTPJ respectively) in ASD and typically developing individuals. A) depicts the raTPJ (yellow) and rpTPJ (green) seed ROIs superimposed on the Oren-Nayer brain surface in <u>Surf</u> <u>Ice</u> viewer. Centre of gravity (in MNI space) for the raTPJ: x = 59, y = -37, z = 30; and for the rpTPJ: x = 54, y = - 55, z = 26. B) and C) depict the group differences in dynamic FC between raTPJ and rpTPJ in the data-driven temporal variability component obtained from the dynamic independent component analysis (dyn-ICA) in the Exploratory and Replication datasets, respectively. D) and E) depict the association and corresponding 95% Confidence Intervals between dyn-ICA and autistic traits, measured with the Social Responsiveness Scale (SRS) total scores in the Exploratory and Replication datasets, respectively.

In the Exploratory dataset, a temporal modulation smoothing kernel of 37.4s (i.e., 22 TRs) was defined in the *Dyn-ICA* tab of the CONN toolbox and the raTPJ and rpTPJ masks were introduced as seeds. In the Replication dataset, the temporal window was set to 39s (i.e., 13 TRs). The dyn-ICA analysis is performed in different steps. First, fMRI data is concatenated across subjects and iterative dual regression is used to estimate the subject-specific matrix of connectivity changes and the subject-specific timeseries of the temporal modulatory component. Next, fastICA is used to obtain the dynamic, data-driven, independent component. Finally, generalised psychophysiological interaction (gPPI) back-projection is implemented to obtain each subject’s dyn-ICA matrix (connectivity values between raTPJ and rpTPJ), with the estimated component as the gPPI psychological factor (57).

### Dynamic FC analysis in the Replication dataset (sliding-window approach)

An ROI-to-ROI sliding-window dynamic FC analysis (41,44) was additionally carried out in the Replication dataset to expand the results obtained with the dyn-ICA approach. ROI-to-ROI dynamic FC analyses return connectivity matrices representing the temporal variability in dynamic FC between pairs of seeds. The window length was set to 13 TRs (39s) with a sliding window step of 1 TR (3s) to ensure smooth transitions between windows, producing 108 temporal windows in total (120 volumes – 13 TRs + 1). These parameters were selected in order to find a balance between the sensitivity and specificity of the window, i.e., to define a window short enough to capture rapidly shifting dynamics of FC but long enough to avoid spurious fluctuations (58, 59).

For the first level analysis, the FC values (Fisher’s z transformed correlation coefficient; z-scores) between the pair of ROIs were computed for each sliding-window and for each subject. This yielded a set of beta-maps per subject that were introduced into a general linear model for ROI-to-ROI group-level analysis. Effects were considered significant if p_uncorrected_<0.01 (two-sided) and p-FDR (False Discovery Rate) _corrected_ <0.05.

In order to investigate the states of connectivity, FC patterns were estimated by classifying each subject’s ROI-to-ROI connectivity values from each of the 108 windows into five states (47): high negative (z-scores < -0.50), moderate negative (−0.5 ≤ z-scores < -0.25), low-uncorrelated (−0.25 ≤ z-scores ≤ 0.25), moderate positive (0.25 < z-scores ≤ 0.50) and high positive (z-scores > 0.5). We then extracted three indices quantifying the connectivity states. These included the proportion of windows that fell into each particular state (Proportion); the average number of continuous windows attributed to the same state (Mean Dwell Time – MDT) (41), and the probability of entering a specific connectivity state (Probability of Transition - PT), which was depicted by the number of times a participant entered that specific state of connectivity over the total number of state changes (40). All indices were computed using in-house scripts (available online at https://github.com/laurabravo97/dynamic_functional_connectivity_analysis) written in Python (https://www.python.org/).

### Static FC analysis

Static FC was also assessed to ascertain whether stationary and dynamic FC analyses produce complementary or overlapping information. For the first-level analysis, the correlation coefficients between the full time series of the raTPJ and the rpTPJ were obtained for each subject. The resulting static beta maps were introduced into a general linear model for ROI-to-ROI group-level analysis and, similar to dynamic FC analyses, effects were considered significant if p_uncorrected_<0.01 and p_FDR corrected_ <0.05.

### Statistical analyses

Two-sample t-test was used to compare demographic characteristics and autistic traits between the ASD and TD groups. Group differences in dyn-ICA and static FC were assessed using independent samples t-test. Pearson’s correlation analyses between dyn-ICA and connectivity state indices (i.e., Proportion, MDT, and PT) were performed in the Replication dataset to further explore the dynamic relationship between raTPJ and rpTPJ functional connectivity. Multiple comparison correction (false discovery rate, FDR) was performed for all connectivity matrices (dyn-ICA, proportion of windows, MDT, and PT) using Benjamini-Hochberg procedure. In addition, three repeated measures ANOVAs were performed in the Replication dataset to explore group differences in proportion of windows, MDT and PT; with “group” as the between subject variable and “state of connectivity” (i.e., high negative, moderate negative, low uncorrelated, moderate positive and high positive) as the within subject variable (48).

In both datasets, the association between autistic traits (quantified with SRS Total scores) and dyn-ICA was examined using Pearson’s correlation analyses for the entire samples. These analyses were followed by General Linear Models (GLM) to explore main effects and interaction effects of dyn-ICA and group in predicting SRS total scores. Similarly, in the Replication dataset Pearson’s correlation analyses were performed to examine the association between SRS total scores and connectivity state indices. Significant results were followed by GLMs to evaluate between-subjects effects of connectivity state indices, group, and their interaction in predicting SRS Total scores. All p-values reported are FDR corrected unless otherwise stated.

## Results

### Participant characteristics

Table 1 summarizes demographic characteristics and autistic traits measured with the SRS in both the Exploratory and Replication datasets. Two-sample T-tests showed no differences between the ASD and TD groups in age or FIQ in either dataset. Across the two datasets, however, the ASD group scored significantly higher on the SRS Total and all subscales (all *ps* < .05).

### Group differences in dynamic FC

Independent samples T-tests revealed significant differences between the ASD and TD groups in the dynamic connectivity between raTPJ and rpTPJ in both the exploratory (*t* = -2.34, *p* = 0.027, Cohen’s d = 0.88, Figure 1B) and replication datasets (*t* = -2.21, *p* = 0.031, Cohen’s d = 0.60, Figure 1C). In line with our prediction, in both datasets, the ASD group showed decreased dynamic resting-state FC (i.e., lower temporal variability) between the anterior and posterior subdivisions of the rTPJ.

### Association between autistic traits and dynamic FC

Pearson correlation was used to assess the association between autistic traits (SRS total scores) and the dynamic FC metric (dyn-ICA) in both the Exploratory and Replication datasets. The analysis revealed a significant negative correlation in the Exploratory (*r* = -0.43, *p* = 0.02) and Replication dataset (*r* = -0.35, *p* = 0.007). Thus, in line with our predictions, increased temporal variability in FC between the raTPJ and rpTPJ was associated with reduced autistic trait expression. To assess for possible group differences in this association, we further conducted regression analyses in the two datasets with dyn-ICA, group, and their interaction as predictors and the SRS total score as the outcome measure. In the Exploratory dataset, the regression yielded a significant model, F (3,25) = 6.26, *p* = 0.003, η_p_ ^2^ = 0.429, but no significant main effects or interaction effects were found, probably due to the small sample size. Subsequently, a main effect only model (F (2,26) = 9.76, *p* = 0.001, η_p_ ^2^ = 0.429) revealed a main effect of group, β(se) = 28.92(8.62), t = 3.35, *p* = 0.02, η_p_ ^2^ = 0.302, such that the ASD group presented significantly higher autistic traits (see Figure 1D). In the Replication dataset, the model (F (3, 53) = 56.30, *p* < 0.001, η_p_ ^2^ = 0.761) revealed a significant interaction between dyn-ICA and group in predicting autistic traits, F (1,53) = 6.08, *p* = 0.017, η_p_ ^2^ = 0.103 (see Figure 1E). Specifically, higher dynamic FC was associated with reduced autistic traits in individuals with ASD, β(se) = -138.51(54.91), t = -2.52, *p* = 0.018, η_p_^2^ = 0.197, but not in the TD controls, *p* = 0.75.

### Specific connectivity states (in Replication dataset)

Repeated measures ANOVAs of the connectivity state indices (Proportion, Mean Dwell Time and Probability of Transition) highlighted a global difference in these metrics as a function of connectivity state (all p_s_ < 0.001) but this did not change based on group and there was no interaction between state and group either (results of the ANOVAs are presented in Supplementary Material Table S1 and graphed together in Figure S1).

To further assess the dynamic connectivity between the raTPJ and rpTPJ and its association with autistic traits, we first performed Pearson correlations across the entire sample between the connectivity state indices and SRS total scores. Pearson correlations revealed that autistic trait expression was *negatively* correlated with Probability of Transition to and Proportion of high negative connectivity states (*r_PT_* = -0.31, *p* = 0.02 and *r_proportion_*= -0.35, *p* = 0.007). This suggests that individuals for which the high negative state (antagonistic relation) was more prominent tended to show lower levels of autistic traits. In addition, autistic trait expression was also *positively* correlated with Probability of Transition to and Proportion of high positive states (*r_PT_* = 0.33, *p* = 0.01 and *r_proportion_* = 0.32, *p* = 0.01). Thus, individuals for which the high positive state was more prominent tended to show higher levels of autistic traits. There were no significant correlations between the Mean Dwell Time of any of the five states and total SRS scores (−0.22 < *rs* < 0.25, all *ps* > 0.057).

Next, we used regression analyses to establish whether these significant associations between connectivity metrics and autistic traits are dependent on group. We, therefore, used group, index of connectivity state, and their interaction to predict autistic trait expression in each of the connectivity states separately (see Supplementary Material Table S2 for full details of the regression models). Results in the high negative connectivity state (see Figures 2A, C) revealed significant interactions between group and Probability of Transition, F(1,53) = 5.29, *p* = 0.025, η_p_ ^2^ = 0.091, and Proportion of windows, F(1,53) = 5.77, *p* = 0.020, η_p_ ^2^ = 0.098. Examining the interaction effects revealed that lower Probability of Transition to (β(se) = -267.20(97.60), t = -2.74, *p* = 0.020, η_p_^2^ = 0.19) and lower Proportion of (β(se) = -129.25(52.37), t = -2.472, *p* = 0.018, η_p_ ^2^ = 0.197) the high negative connectivity state were associated with increased autistic trait expression in ASD participants, but not in TD controls (*p*_PT_ = 0.58 and *p*_proportion_ = 0.72). In the high positive connectivity state (Figure 2B, D), the models revealed only a significant main effect for group (Probability of transition: F(1,53)=12.18, p = 0.001, η_p_^2^ = 0.187; Proportion of windows: F(1,53)=20.82, p < 0.001, η ^2^ = 0.282), such that autistic trait expression was higher in the ASD than in the TD group. Thus, it appears that prominence of the highly negative state is associated with reduced autistic traits, and that this is particularly pronounced in the ASD group.

**Figure 2.**
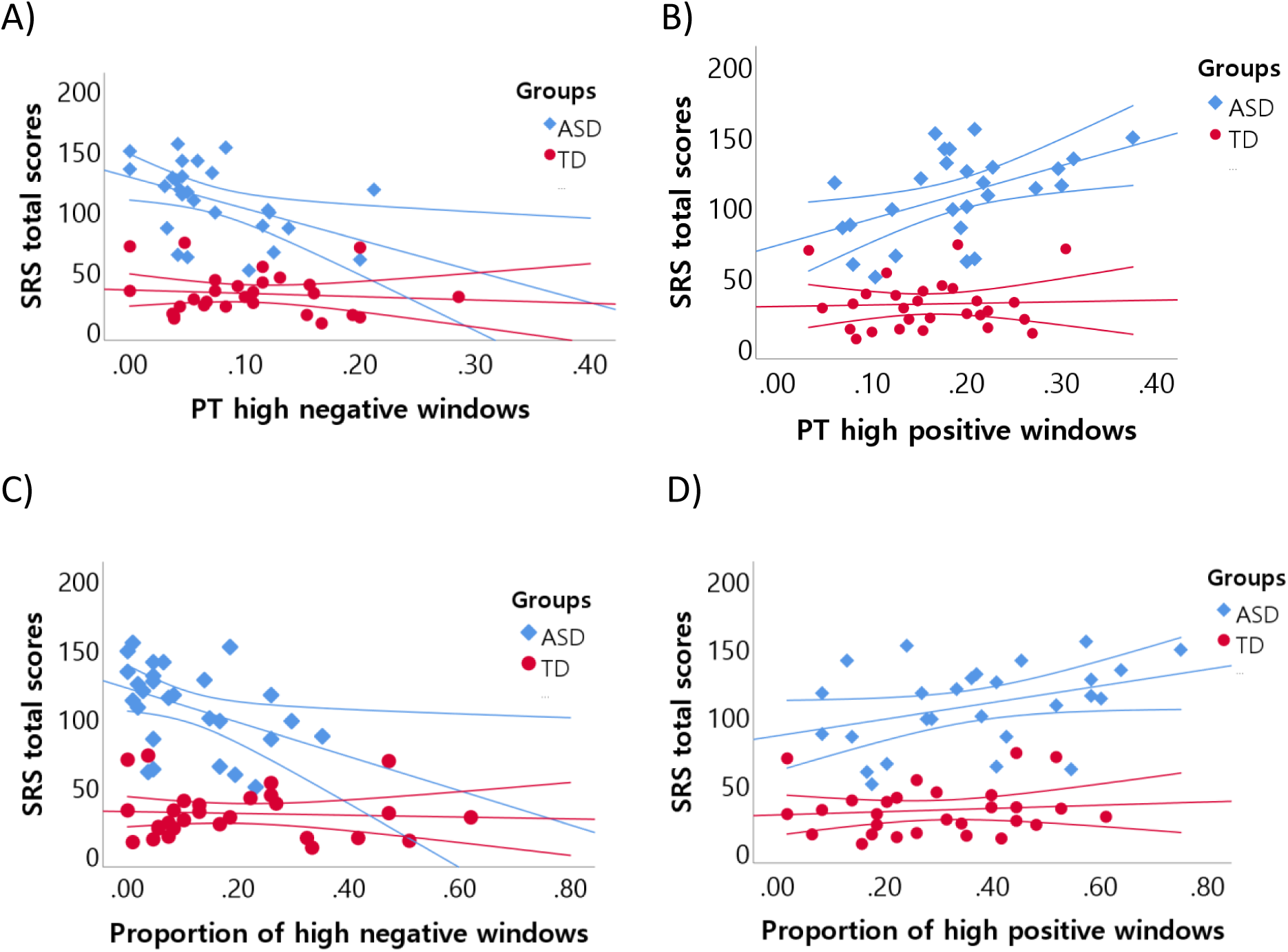
Association between temporal variability metrics (probability of transition,PT, and proportion of windows) and autistic trait expression in both high negative and high positive connectivity states in the ASD and the typically developing groups of the Replication dataset. Autistic trait expression is quantified with the total score of the Social Responsiveness Scale (SRS). A) depicts the association of PT with SRS total scores in the high negative connectivity state; B) depicts the association of PT with SRS total scores in the high positive connectivity state; C) depicts the association of proportion of windows with SRS total scores in the high negative connectivity state; and D) depicts the association of proportion of windows with SRS total scores in the high positive connectivity state. All regression lines are depicted with the corresponding 95% Confidence Intervals.

Finally, we assessed the possible link between the dyn-ICA measure representing the global temporal variability in functional connectivity and the specific states of connectivity (from high negative to high positive). To this end, we assessed the Pearson correlations between dyn-ICA and the three indices of the five connectivity states. The analysis revealed significant *positive* correlation between dyn-ICA and the probability of transitioning to a high negative connectivity state (*r* = 0.43, *p_FDR_*= 0.003), suggesting that increased global temporal variability was associated with a stronger tendency for the raTPJ-rpTPJ connectivity to move towards a high negatively correlated (antagonistic) state. In addition, we observed a significant *negative* correlation between dyn-ICA and the probability of transitioning to a high positive connectivity state (*r* = -0.42, *p_FDR_*= 0.003). Thus, increased global temporal variability was also associated with a decreased tendency for the raTPJ-rpTPJ connectivity to shift towards a high positively correlated state (see Supplementary Material Table S3 for full details of the correlations between dyn-ICA and connectivity state indices).

### Static functional connectivity

To assess whether static functional connectivity could also be useful here, we repeated the same analyses we report with the dyn-ICA metric with the static functional connectivity between the raTPJ and the rpTPJ. We observed no significant differences in static FC between the ASD and TD groups in either the Exploratory (*t* = 0.71, *p* = 0.482) or the Replication (*t* = 1.37, *p* = 0.176) datasets. There were also no significant correlations between static FC and SRS total scores in either the Exploratory (*r* = 0.11, *p* = 0.568) or the replication (*r* = 0.25, *p* = 0.064) datasets.

## Discussion

The present study is the first to explore the dynamic FC between the anterior and posterior subdivisions of the rTPJ in ASD and neurotypical adults as a window to understanding the relationship between attention reorienting and ToM in ASD. For this purpose, the dynamic ICA (57) and the sliding-window (41, 42) approaches were used in two datasets (Exploratory and Replication) to yield specific metrics of dynamic functional connectivity both on the global level (dyn-ICA – for both the Exploratory and Replication datasets) and on specific connectivity state level (proportion of windows, Mean Dwell Time and Probability of Transition – only in the Replication dataset). Consistent with our predictions, the results in the Exploratory dataset highlighted a decreased dynamic FC between the posterior and anterior divisions of the right TPJ in the ASD group. This finding was then corroborated in the Replication dataset, with similar reduction in global dynamic FC in ASD compared to neurotypical controls. Moreover, especially in the replication dataset, this global signal of dynamic functional connectivity was associated with the autistic trait expression in ASD so that reduced dynamic functional connectivity was associated with higher autistic traits.

The reduced dyn-ICA signal in ASD we report here fits with similar findings regarding dynamic FC in this population (although previously a whole-brain analysis has been the most common approach). For example, Yao et al. (49) performed time-varying FC analyses in ASD and found reduced dynamic connectivity between the posterior (precuneus/posterior cingulate gyrus) and anterior (medial superior frontal gyrus) hubs of the default mode network, which similarly to what we report was also associated with social difficulties. This is consistent with the general idea that a more variable connectivity between regions contribute to enhanced cognitive processing (41) and would offer greater flexibility to neural networks to jointly function when a task is presented (60). Moreover, if variable or dynamic connectivity between brain regions is particularly important in integrative functions (53), the decreased dynamic FC between raTPJ (part of the attention reorienting network) and rpTPJ (part of the ToM network) shown here by the ASD group may point to the importance of integrative attention and ToM processes in driving successful social interaction. It is therefore plausible that at least some of the difficulty in social interaction in ASD might be explained by such reduced integration of the attention and ToM processes. It is not clear, however, if this is a consequence of separately or independently impaired attention or ToM in ASD (e.g., the precursory nature of attention shifting problems in ASD) (28) or that the integration itself is the source of the social interaction impairment irrespective of the attention and ToM processes per se.

Our sliding window approach, applied in the Replication dataset enabled us to identify whether specific states of connectivity play an important role through the categorization of connectivity states into five different categories (from highly negatively correlated to highly positively correlated). Previous research has suggested that, at least in static FC, there appears to be an antagonistic relationship between attention and mentalizing networks (36) and between the raTPJ and rpTPJ specifically (30). In the context of the sliding window approach, this would translate to a high proportion of (and possibly higher rates of transition to and dwell time in) a high negative connectivity state. The ANOVA comparing these metrics across the ASD and neurotypicals in our study did not reveal a significant group difference. Critically, however, we found clear association in the ASD group between proportion and probability of transition into the high negative connectivity state and autistic trait expression. Reduced prominence of the highly negative state was associated with greater expression of autistic traits. This was also complimented (although in a less pronounced way) by the association between increased prominence of the highly positive state and higher autistic traits. Thus, it appears that in ASD, participants for whom the highly negative state is less pronounced than the highly positive state are those who show reduced social responsiveness.

A few previous studies utilising different versions of a sliding window approach with a whole brain analysis have identified not only reduced functional connectivity but also a tendency in ASD to spend more time in a weak FC state (or a state of underconnectivity) compared to TD individuals (45,49). Rabany and colleagues (48), for instance, compared the temporal variability and connectivity patterns between 56 independent components in individuals with ASD, schizophrenia and typically developing and reported a generally decreased dynamic FC in the ASD population due to an increased dwell time in a state of underconnectivity. While we did not find evidence for such a group difference in an uncorrelated state in our study, it is worth noting that our focus on the raTPJ and rpTPJ meant the most indicative correlation states were the highly negative and highly positive states. The present results showed that individuals spent significantly more time in both highly negative and highly positive connectivity states compared to an uncorrelated state (see Figure S1-B and Table S4 for pairwise comparisons). In addition, our findings pointed to individual differences in the way these states may contribute to behavioural traits in ASD rather than a whole group difference in these metrics (although a non-significant tendency towards greater proportion of windows in the high positive connectivity state was seen in the ASD group; see Figure S1-A).

The relevance of the highly negative and highly positive correlation states was also exemplified through their association with global dynamic FC. Correlation analyses revealed that increased dynamic FC was associated with greater probability of transition to the high negative connectivity state but also reduced probability of transition to the high positive connectivity state. Generally, across the various FC state indices, when moving from highly negative to highly positive states (through the intermediary connectivity states) there is a gradual shift from positively correlated to negatively correlated dynamic FC (see Table S3). Thus, dynamic FC between the raTPJ and rpTPJ seems to be associated with a balance between highly negative and highly positive states. However, when participants present with a shift away from this balance (away from the negative state and towards the positive state), they seem to also show reduced dynamic FC, as well as increased autistic trait expression.

Our exclusive focus on the rTPJ given its relevance as a possible interaction point between attention reorienting and ToM (27, 28) has enabled us to identify global dynamic FC indices as well as specific meaningful connectivity states. However, as noted above, it is harder to compare our findings to previous attempts at identifying dynamic FC metrics in ASD. Although the field of dynamic FC is rapidly growing, much remains unknown about the methodology and neurocognitive implications of this analysis (61). The lack of a grounded theory regarding the parameters for the sliding window technique for instance (especially window length and sliding step) makes it harder to compare findings across different investigations. In spite of this, the present study confirmed the importance of investigating the FC between interrelated neural networks from a time-varying perspective. Indeed, while we report significant differences (both on a group and on an individual levels) using dynamic FC metrics, using a static FC metric did not yield significant findings. Moreover, previous static FC investigations indicated an anticorrelated connectivity between raTPJ and rpTPJ (30), but dynamic FC analyses have revealed that the relationship between these regions is not so straightforward and that high positive states also play an important role. It is therefore possible that inconsistencies in previous research looking at FC in ASD relate to the limitation of the static FC approach to capture different connectivity states present throughout the scan.

In conclusion, the present study suggests that difficulties in social interaction in autistic individuals could be linked to an atypical interaction between the ToM and attention orienting networks, expressed as reduced dynamic functional connectivity between the anterior and posterior subdivisions of the right TPJ. We suspect that this reduced decreased dynamic connectivity hinders the integrative process of ToM and attention orienting, which combined contribute to the manifestation of social difficulties (62). It appears that the reduced dynamic FC is associated with an imbalance between positive and negative correlation states. Importantly, both the reduced dynamic FC and the reduced prominence of a highly negative connectivity state are predictive of increased autistic trait expression. Our findings provide strong support for the import of assessing dynamic FC in ASD and the sensitivity of this measure to individual differences in social abilities.

## Acknowledgments and Disclosures

The authors declare that they have no conflict of interest. All authors have seen and agree with the content of the manuscript and there is no financial interest to report.

## References

1. American Psychiatric Association (2013). Diagnostic and Statistical Manual of Mental Disorders. 5th edition.

2. Keehn, B., Lincoln, A., Müller, R. A., & Townsend, J. (2010). Attentional networks in children and adolescents with autism spectrum disorder. Journal of Child Psychology and Psychiatry, 51(11), 1251–1259. https://doi.org/10.1111/j.1469-7610.2010.02257.x

3. Lai, M.C., Lombardo, M.V., & Baron-Cohen, S. (2014). Autism. Lancet, 383, 896–910. https://doi.org/10.1016/S0140-6736(13)61539-1

4. Leekam S. (2016). Social cognitive impairment and autism: what are we trying to explain? Philosophical Transactions of the Royal Society B Biological Sciences, 371(1686): 20150082. https://doi.org/10.1098/rstb.2015.0082

5. Spaniol, M. M., Magalhães, J., Mevorach, C., Shalev, L., Teixeira, M. C. T. V., Lowenthal, R., & de Paula, C. S. (2021). Association between attention, nonverbal intelligence and school performance of school-age children with Autism Spectrum Disorder from a public health context in Brazil. Research in developmental disabilities, 116, 104041. https://doi.org/10.1016/j.ridd.2021.104041

6. Baron-Cohen, S., Leslie, A. M. & Frith, U. (1985). Does the autistic child have a “theory of mind”? Cognition, 21, 37–46. https://doi.org/10.1016/0010-0277(85)90022-8

7. Hill, E.L., Frith, U., (2003). Understanding autism: insights from mind and brain. Philosophical Transactions of the Royal Society of London B: Biological Sciences, 358, 281–289. https://doi.org/10.1098/rstb.2002.1209

8. Jones, C.R., Simonoff, E., Baird, G., Pickles, A., Marsden, A.J., Tregay, J., Happé, F. & Charman, T., (2018). The association between theory of mind, executive function, and the symptoms of autism spectrum disorder. Autism Research, 11(1), 95–109. https://doi.org/10.1002/aur.1873

9. Abu-Akel, A. and Bailey, A.L. (2001). Indexical and symbolic referencing: what role do they play in children’s success on theory of mind tasks? Cognition, 80(3), 263–81. https://doi.org/10.1016/s0010-0277(00)00149-9

10. Wimmer, H. and Perner, J. (1983). Beliefs about beliefs: representation and constraining function of wrong beliefs in young children’s understanding of deception. Cognition, 13, 103–128. https://doi.org/10.1016/0010-0277(83)90004-5

11. Castelli F., Frith C., Happe F., Frith U. (2002). Autism, Asperger syndrome and brain mechanisms for the attribution of mental states to animated shapes. Brain, 125(Pt 8), 1839–1849. https://doi.org/10.1093/brain/awf189

12. Cole, E.J., Barraclough, N.E. & Andrews, T.J. (2019). Reduced connectivity between mentalizing and mirror systems in autism spectrum condition. Neuropsychologia, 122, 88–97. https://doi.org/10.1016/j.neuropsychologia.2018.11.008

13. Dichter G. S. (2012). Functional magnetic resonance imaging of autism spectrum disorders. Dialogues in clinical neuroscience, 14(3), 319–351. https://doi.org/10.31887/DCNS.2012.14.3/gdichter

14. Kana, R.K., Maximo, J.O., Williams, D.L., Keller, T.A., Schipul, S.E., Cherkassky, V.L., Minshew, N.J. and Just, M.A., (2015). Aberrant functioning of the theory-of-mind network in children and adolescents with autism. Molecular autism, 6(1), 1–12. https://doi.org/10.1186/s13229-015-0052-x

15. Keehn, B., Müller, R. A., & Townsend, J. (2013). Atypical attentional networks and the emergence of autism. Neuroscience and Biobehavioral Reviews, 37(2), 164–183. https://doi.org/10.1016/j.neubiorev.2012.11.014

16. Abu-Akel, A., Apperly, I., Spaniol, M. M., Geng, J. J., & Mevorach, C. (2018). Diametric effects of autism tendencies and psychosis proneness on attention control irrespective of task demands. Scientific reports, 8(1), 8478. https://doi.org/10.1038/s41598-018-26821-7

17. Posner MI, Snyder CRR, Davidson BJ (1980) Attention and the detection of signals. J Exp Psychol Gen, 109, 160–174. https://doi.org/10.1037//0096-3445.109.2.160

18. Dawson, G., Meltzoff, A.N., Osterling, J., Rinaldi, J., & Brown, E. (1998). Children with autism fail to orient to naturally occurring social stimuli. Journal of Autism and Developmental Disorders, 28, 479–485. https://doi.org/10.1023/A:1026043926488

19. Dawson, G., Toth, K., Abbott, R., Osterling, J., Munson, J., Estes, A., & Liaw, J. (2004). Early social attention impairments in autism: Social orienting, joint attention, and attention to distress. Developmental Psychology, 40, 271–283. https://doi.org/10.1037/0012-1649.40.2.271

20. Greene, D.J., Colich, N., Iacoboni, M., Zaidel, E., Bookheimer, S.Y. & Dapreto, M. (2011). Atypical neural networks for social orienting in autism spectrum disorders. Neuroimage, 56, 354–362. https://doi.org/10.1016/j.neuroimage.2011.02.031

21. Gomot M, Bernard FA, Davis MH, Belmonte MK, Ashwin C, Bullmore ET, Baron-Cohen S. (2006). Change detection in children with autism: An auditory event-related fMRI study. Neuroimage, 29:475–484. https://doi.org/10.1016/j.neuroimage.2005.07.027

22. Gomot M, Belmonte MK, Bullmore ET, Bernard FA, Baron-Cohen S. (2008). Brain hyper-reactivity to auditory novel targets in children with high-functioning autism. Brain, 131, 2479–2488. https://doi.org/10.1093/brain/awn172

23. Mundy, P., & Newell, L. (2007). Attention, joint attention, and social cognition. Current Directions in Psychological Science, 16, 269–274. https://doi.org/10.1111/j.1467-8721.2007.00518.x

24. Corbetta, M., Patel, G., and Shulman, G. (2008). The reorienting system of the human brain: From environment to theory of mind. Neuron, 58, 306–324. https://doi.org/10.1016/j.neuron.2008.04.017

25. Decety, J. & Lamm, C. (2007). The role of the right temporoparietal junction in social interaction: how low-level computational processes contribute to meta-cognition. Neuroscientist, 13, 580–593. https://doi.org/10.1177/1073858407304654

26. Zwaigenbaum, S. Bryson, T. Rogers, W. Roberts, J. Brian, P. Szatmari. (2005). Behavioral manifestations of autism in the first year of life. International Journal of Developmental Neuroscience, 23, 143–152. https://doi.org/10.1016/j.ijdevneu.2004.05.001

27. Devaney, K. J. (2018). Functional MRI investigations of human temporoparietal junction: attention, response inhibition, theory of mind, and long-term meditation effects (Doctoral thesis, Boston University, Boston, United States). Retrieved from https://hdl.handle.net/2144/31877

28. Kubit, B. & Jack, A. I. (2013). Rethinking the role of the rTPJ in attention and social cognition in light of the opposing domains hypothesis: Findings from an ALE-based meta-analysis and resting-state functional connectivity. Frontiers in Human neuroscience, 7 (323), 1–18. https://doi.org/10.3389/fnhum.2013.00323

29. Abu-Akel, A., Apperly, I. A., Wood, S. J. & Hansen, P. C. (2017). Autism and psychosis expressions diametrically modulate the right temporoparietal junction. Social Neuroscience, 12(5), 506–518. https://doi.org/10.1080/17470919.2016.1190786

30. Bzdok, D., Langner, R., Schilbach, L., Jakobs, O., Roski, C., Caspers, S., Laird, A. R., Fox, P. T., Zilles, K., & Eickhoff, S. B. (2013). Characterization of the temporo-parietal junction by combining data-driven parcellation, complementary connectivity analyses, and functional decoding. NeuroImage, 81, 381–392. https://doi.org/10.1016/j.neuroimage.2013.05.046

31. Mars, R. B., Sallet, J., Schüffelgen, U., Jbabdi, S., Toni, I., & Rushworth, M. F. (2012). Connectivity-based subdivisions of the human right “temporoparietal junction area”: evidence for different areas participating in different cortical networks. Cerebral cortex (New York, N.Y. : 1991), 22(8), 1894–1903. https://doi.org/10.1093/cercor/bhr268

32. Schuwerk, T., Schurz, M., Müller, F., Rupprecht, R. & Sommer, M. (2017). The rTPJ’s overarching cognitive function in networks for attention and theory of mind. Social Cognitive and Affective Neuroscience, 12(1), 157–168. https://doi.org/10.1093/scan/nsw163

33. Krall, S.C., Volz, L.J., Oberwelland, E., Grefkes, C., Fink, G.R., Konrad, K. (2016). The right temporoparietal junction in attention and social interaction: a transcranial magnetic stimulation study. Human Brain Mapping, 37(2), 796–807. https://doi.org/10.1002/hbm.23068

34. Krall, S.C., Rottschy, E., Oberwelland, E., Bzdok, D., Fox, P. T., Eickhoff, S. B., Fink, G. R. & Konrad, K. (2015). The role of the right temporoparietal junction in attention and social interaction as revealed by ALE meta-analysis. Brain Struct. Funct., 220, 587–604. https://doi.org/10.1007/s00429-014-0803-z

35. Jack, A. I., Dawson, A. J., Begany, K. L., Leckie, R. L., Barry, K. P., Ciccia, A. H., et al. (2012). fMRI reveals reciprocal inhibition between social and physical cognitive domains. Neuroimage 66C, 385–401. https://doi.org/10.1016/j.neuroimage.2012.10.061

36. Fox, M. D., Snyder, A. Z., Vincent, J. L., Corbetta, M., Van Essen, D. C., and Raichle, M. E. (2005). The human brain is intrinsically organized into dynamic, anticorrelated functional networks. Proc. Natl. Acad. Sci. U.S.A., 102, 9673–9678. https://doi.org/10.1073/pnas.0504136102

37. Alaerts, K., Woolley, D. G., Steyaert, J., Di Martino, A., Swinnen, S. P., & Wenderoth, N. (2014). Underconnectivity of the superior temporal sulcus predicts emotion recognition deficits in autism. Social cognitive and affective neuroscience, 9(10), 1589–1600. https://doi.org/10.1093/scan/nst156

38. von dem Hagen, E.A., Stoyanova, R.S., Baron-Cohen, S., & Calder, A.J. (2013). Reduced functional connectivity within and between ‘social’ resting state networks in autism spectrum conditions. Social Cognitive and Affective Neuroscience, 8, 694–701. https://doi.org/10.1093/scan/nss053

39. Chien, H. Y., Lin, H. Y., Lai, M. C., Gau, S. S. F. & Tseng, W. Y. I. (2015). Hyperconnectivity of the right temporo-parietal junction predicts social difficulties in boys with Autism Spectrum Disorder. Autism Research, 8, 427–441. https://doi.org/10.1002/aur.1457

40. Li, Y., Zhu, Y., Nguchu, B. A., Wang, Y., Wang, H., Qiu, B. and Wang, X. (2020). Dynamic Functional Connectivity Reveals Abnormal Variability and Hyper-connected Pattern in Autism Spectrum Disorder. Autism Research, 13, 230–243. https://doi.org/10.1002/aur.2212

41. Allen, E. A., Damaraju, E., Plis, S. M., Erhardt, E. B., Eichele, T., & Calhoun, V. D. (2014). Tracking whole brain connectivity dynamics in the resting state. Cerebral Cortex, 24(3), 663–676. https://doi.org/10.1093/cercor/bhs352

42. Calhoun, V. D., Miller, R., Pearlson, G. and Adali, T. (2014). The Chronnectome: time-varying connectivity networks as the next frontier in fMRI data discovery. Neuron, 84, 262–274. https://doi.org/10.1016/j.neuron.2014.10.015

43. Hutchison, R. M., Womelsdorf, T., Gati, J. S., Everling, S., Menon, R. S. (2013a). Resting-state networks show dynamic functional connectivity in awake humans and anesthetized macaques. Human Brain Mapping, 34, 2154–2177. https://doi.org/10.1002/hbm.22058

44. Hutchison, R. M., Womelsdorf, T., Allen, E. A., Bandettini, P. A., Calhoun, V. D., Corbetta, M. et al (2013b). Dynamic functional connectivity: promise, issues, and interpretations. Neuroimage, 80, 360–378. https://doi.org/10.1016/j.neuroimage.2013.05.079

45. Chen, H., Nomi, J. S., Uddin, L. Q., Duan, X. J., & Chen, H. F. (2017). Intrinsic functional connectivity variance and state speciFic under-connectivity in autism. Human Brain Mapping, 38(11), 5740–5755. https://doi.org/10.1002/hbm.23764

46. Rashid, B., Blanken, L. M. E., Muetzel, R. L., Miller, R., Damaraju, E., Arbabshirani, M.R., Calhoun, V. et al. (2018). Connectivity dynamics in typical development and its relationship to autistic traits and autism spectrum disorder. Human Brain Mapping, 39(8), 3127–3142. https://doi.org/10.1002/hbm.24064

47. Kaiser, R. H., Whitfield-Gabrieli, S., Dillon, D. G., Goer, F., Beltzer M., Minkel, J., Smoski, M., Dichter, and G., Pizzagalli, D. A. (2016). Dynamic resting-state functional connectivity in major depression. Neuropsychopharmacology, 41, 1822–1830. https://doi.org/10.1038/npp.2015.352

48. Rabany, L., Brocke, S., Calhoun, V. D., Pittman, B., Corbera, S., Hyatt, C. J., Wexler, B., Bell, M. D., Pelphrey, K., Pearlson, G. D. & Assaf, M. (2019). Dynamic functional connectivity in schizophrenia and autism spectrum disorder: Convergence, divergence, and classification. NeuroImage, 24. https://doi.org/10.1016/j.nicl.2019.101966

49. Yao, Z. J., Hu, B., Xie, Y. W., Zheng, F., Liu, G. Y., Chen, X. J., & Zheng, W. H. (2016). Resting-state time-varying analysis reveals aberrant variations of functional connectivity in autism. Frontiers in Human Neuroscience, 10, 463. https://doi.org/10.3389/fnhum.2016.00463

50. Di Martino, A., Yan, C.G., Li, Q., Denio, E., Castellanos, F.X., Alaerts, K., et al. (2014). The autism brain imaging data exchange: Towards a large-scale evaluation of the intrinsic brain architecture in autism. Molecular Psychiatry, 19, 659– 667. https://doi.org/10.1038/mp.2013.78

51. Di Martino, O’Connor D., Chen, B., et al. (2017). Enhancing studies of the connectome in autism using the autism brain imaging data exchange II. Scientific Data, 4, 170010. https://doi.org/10.1038/sdata.2017.10

52. Hellyer, P. J., Scott, G., Shanahan, M., Sharp, D. J. & Leech, R. (2015). Cognitive flexibility through metastable neural dynamics is disrupted by damage to the structural connectome. J. Neurosci., 35, 9050–9063. https://doi.org/10.1523/JNEUROSCI.4648-14.2015

53. Shine, J. M. et al. (2016). The dynamics of functional brain networks: integrated network states during cognitive task performance. Neuron, 92, 544–554. https://doi.org/10.1016/j.neuron.2016.09.018

54. Abu-Akel, A., Allison, C., Baron-Cohen, S., Heinke, D. (2019). The distribution of autistic traits across the autism spectrum: Evidence for discontinuous subpopulations underlying the autistic continuum. Molecular Autism, 10(24). https://doi.org/10.1186/s13229-019-0275-3

55. Constantino, J.N., Gruber, C.P. (2005). Social Responsiveness Scale. Los Angeles, CA: Western Psychological Services.

56. Whitfield-Gabrieli, S., Nieto-Castanon, A. (2012). Conn: a functional connectivity toolbox for correlated and anticorrelated brain networks. 2, 125–141. https://doi.org/10.1089/brain.2012.0073

57. Nieto-Castanon, A. (2020). Handbook of functional connectivity Magnetic Resonance Imaging methods in CONN. Boston, MA: Hilbert Press. Retrieved from https://www.researchgate.net/publication/339460691_Handbook_of_functional_connectivity_Magnetic_Resonance_Imaging_methods_in_CONN

58. Leonardi, N. & Van De Ville, D. (2015). On spurious and real fluctuations of dynamic functional connectivity during rest. NeuroImage, 104, 430–436. https://doi.org/10.1016/j.neuroimage.2014.09.007

59. Preti, M. G., Bolton, T. A. W., & Van De Ville, D. (2017). The dynamic functional connectome: State-of-the-art and perspectives. NeuroImage: Clinical, 160, 41–54. https://doi.org/10.1016/j.neuroimage.2016.12.061

60. Nguyen, T. T., Kovacevic, S., Dev, S. I., Lu, K., Liu, T. T., & Eyler, L. T. (2017). Dynamic functional connectivity in bipolar disorder is associated with executive function and processing speed: A preliminary study. Neuropsychology, 31(1), 73–83. https://doi.org/10.1037/neu0000317

61. Chen, J. E., Rubinov, M., & Chang, C. (2017). Methods and Considerations for Dynamic Analysis of Functional MR Imaging Data. Neuroimaging clinics of North America, 27(4), 547–560. https://doi.org/10.1016/j.nic.2017.06.009

62. Ramot, M., Walsh, C., Reimann, G.E. and Martin, A. (2020). Distinct neural mechanisms of social orienting and mentalizing revealed by independent measures of neural and eye movement typicality. Communications Biology, 3(48). https://doi.org/10.1038/s42003-020-0771-1

