## Supplemental Information for "Dynamic functional connectivity in the right temporoparietal junction captures variations in autistic trait expression"

**Running title:** Dynamic FC in right TPJ and autistic trait expression

**Authors**: Laura Bravo Balsa^1^, Ahmad Abu-Akel^2,3^, Carmel Mevorach^4,5,6^

**Affiliations:**

^1^ Department of Forensic and Neurodevelopmental Sciences, Institute of Psychiatry, Psychology & Neuroscience, King’s College London.

^2^ School of Psychological of Sciences, University of Haifa, Haifa, Israel

^3^ Haifa Brain and Behavior Hub, University of Haifa, Haifa, Israel

^4^ School of Psychology, University of Birmingham, Edgbaston, UK

^5^ Centre for Human Brain Health, University of Birmingham, Edgbaston, UK

^6^ Centre for Developmental Science, University of Birmingham, Edgbaston , UK

**Corresponding authors:**

Laura Bravo Balsa

Address: PO Box 23, Dept of Forensic and Neurodevelopmental Sciences, IoPPN, King’s College London, 16 de Crespigny Park Denmark Hill London SE5 8AF

Carmel Mevorach

Address: Centre for Human Brain Health, School of Psychology, University of Birmingham, Edgbaston, Birmingham, B15 2TT, UK.

### **Supplemental Information**

**Data acquisition parameters for the Exploratory dataset (ABIDE I repository: University of Leuven Sample 1).** Anatomical and functional scans were performed on a 3.0 Tesla Philips MR scanner (Best, The Netherlands). A high-resolution T1-weighted structural volume was acquired first in the scanning session (slice thickness = 1.2mm; repetition time (TR) = 9.6ms; echo time (TE) = 4.6ms; matrix size = 256x256; field of view (FOV) = 250x250mm^2^; acquisition time = 6 min 38 s). A resting state scan followed, in which participants were instructed to fixate their eyes on a white cross projected on a black background. The resting state fMRI scans were collected using an echo-planar imaging (EPI) sequence (TR = 1700ms, TE = 33ms; matrix size = 64x64, FOV = 230mm; flip angle 90°; slice thickness = 4mm; number of slices = 32; slice scan order = ascend). The scan lasted for 7 min and comprised of 250 volumes.

**Data acquisition parameters for the Replication dataset (ABIDE II repository: Barrow Neurological Institute).** Anatomical and functional MRI scans were performed on a 3.0 Tesla Philips Ingenia scanner at Barrow Neurological Institute. The scanning session started with an eyes-closed resting functional MRI scans which were collected using an echo-planar imaging (EPI) sequence (TR = 3000ms, TE = 25ms; matrix size = 64x64, FOV = 240x240mm; flip angle 80°; slice thickness = 4mm; number of slices = 50; slice scan order = ascend). The resting state scans lasted 6 minutes and comprised of 120 volumes. The high-resolution T1-weighted structural scans followed the resting scans (slice thickness = 1.2mm; repetition time (TR) = “shortest”; echo time (TE) = “shortest”; matrix size = 244x227; field of view (FOV) = 270x252mm; acquisition time = 5 min 34 s).

**Preprocessing steps (both datasets).** The functional images were: a) realigned to the first volume, b) unwarped to remove distortion-by movement interactions and to estimate and correct the field inhomogeneity inside the scanner, c) slice-timing corrected by temporal interpolation to the middle slice, d) co-registered with structural data, e) spatially normalised into standard Montreal Neurological Institute (MNI) space, f) segmented into white matter, grey matter and cerebrospinal fluid (CSF), and g) spatially smoothed with an isotropic 5mm full-width-at-half-maximum (FWHM) Gaussian kernel (Alaerts et al. 2014).

**Denoising steps (both datasets).** Motion correction was performed using the Artifact Detection Tool (ART, [www.nitrc.org/pro jects/artifact_detect/](http://www.nitrc.org/pro%20jects/artifact_detect/)) to identify outlier scans for scrubbing. In addition, physiological and other sources of noise were identified using CompCor, a method that extracts principal components from the white matter and CSF time courses (Behzadi, Restom, Liau & Liu, 2007). In sum, outlier scans, components from CompCor correction and the six head motion parameters from spatial motion correction were included as confounds in a first-level regression model, followed by linear detrending, despiking and band-pass filtering (0.01-0.08 Hz) to remove low frequency drifts and high frequency noise related to cardiac and respiratory activity (Díez-Cirarda et al. 2018; Kaiser et al., 2016; Wylie, Tregellas, Bear and Legget, 2020).

**Table S1.** ANOVAs repeated measures to explore group differences in connectivity state indices (Proportion of windows, MDT and PT) of the replication sample.

| **Main effects** | **Proportion of windows** | **MDT** | **PT** |
| --- | --- | --- | --- |
| State  (within-subjects) | F(1,56) = 33.79  *p* = 0.000  η_p_^2^ = 0.376 | F(1,56) = 49.75  *p* = 0.000  η_p_^2^ = 0.470 | F(1,56) = 56.57  *p* = 0.000  η_p_^2^ = 0.503 |
| Group  (between-subjects) | F(1,56) = 0  *p = 1*  η_p_^2^ = 0 | F(1,56) = 0.26  *p* = 0.613  η_p_^2^ = 0.005 | F(1,56) = 0.74  *p* = 0.393  η_p_^2^ = 0.013 |
| State * Group  (interaction) | F(1,56) = 2.42  *p* = 0.125  η_p_^2^ = 0.041 | F(1,56) = 1.57  *p* = 0.215  η_p_^2^ = 0.027 | F(1,56) = 1.380  *p* = 0.245  η_p_^2^ = 0.024 |

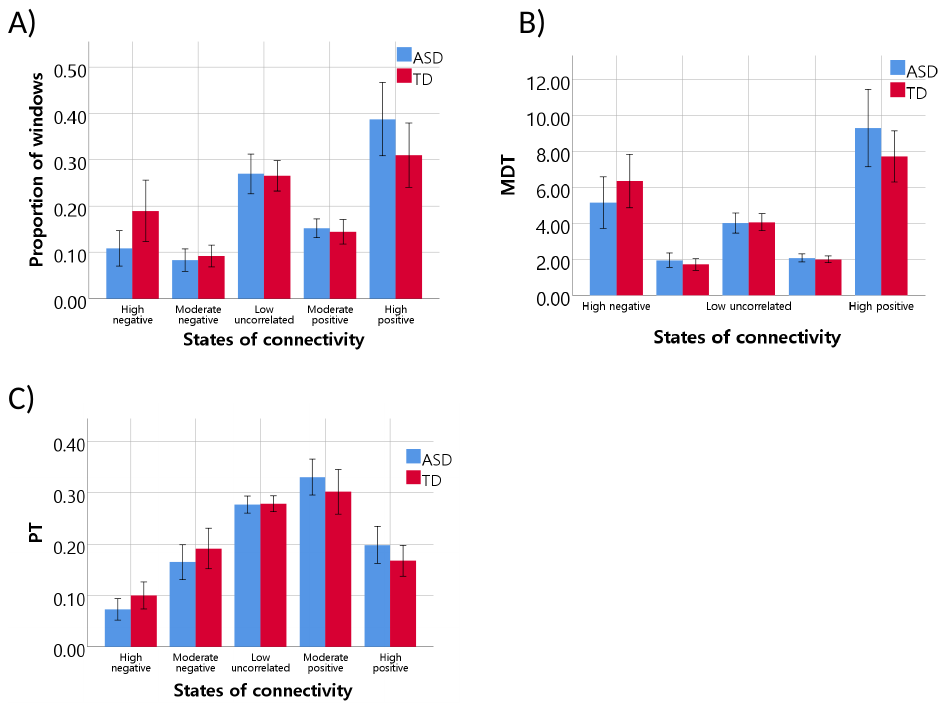

**Figure S1.** Connectivity state indices by group. A) Proportion of windows that fell into each range of connectivity, B) Mean Dwell Time (MDT), i.e., number of consecutive windows (39 seconds each) attributed to the same state, and C) Probability of transition (PT) from other states to a particular one. Both groups spent more time (i.e., increased MDT) in highly negatively and highly positively correlated FC (all p_s_ < 0.005 in pairwise comparisons) and fluctuated more across other states.

**Table S2.** Regression results for the association of connectivity state indices (i.e., proportion of windows and probability of transition, PT), group (ASD and TD) and their interaction with autistic trait expression (SRS total scores) of the replication sample.

| **Predictors** | **Adjusted R^2^** | **ANOVA** | ***p*** | **β** | ***t*** | ***p*** | **Partial eta squared** |
| --- | --- | --- | --- | --- | --- | --- | --- |
| Corrected Model:  PT High Neg. | 0.754 | F(3,53) = 58.25 | <0.001 |  |  |  |  |
| Group |  | F(1,53) = 79.64 | <0.001 | 94.40 | 8.92 | 0.000 | 0.600 |
| PT High Neg. |  | F(1,53) = 8.04 | 0.006 | -27.87 | -0.44 | 0.662 | 0.004 |
| **Group * PT High Neg.** |  | **F(1,53) = 5.29** | **0.025** | **-239.34** | **-2.30** | **0.025** | **0.091** |
| Corrected Model:  PT High Pos. | 0.755 | F(3,53) = 58.47 | <0.001 |  |  |  |  |
| Group |  | F(1,53) = 12.18 | 0.001 | 50.05 | 3.49 | 0.001 | 0.187 |
| PT High Pos. |  | F(1,53) = 5.21 | 0.026 | 11.97 | 0.22 | 0.827 | 0.001 |
| Group * PT High Pos. |  | F(1,53) = 3.79 | 0.057 | 138.95 | 1.95 | 0.057 | 0.067 |
| Corrected Model:  Prop. High Neg. | 0.746 | F(3,53) = 55.59 | <0.001 |  |  |  |  |
| Group |  | F(1,53) = 98.30 | <0.001 | 90.67 | 9.92 | <0.001 | 0.650 |
| Prop. High Neg. |  | F(1,53) = 7.18 | 0.010 | -7.04 | -0.28 | 0.783 | 0.001 |
| **Group * Prop. High Neg.** |  | **F(1,53) = 5.77** | **0.020** | **-122.21** | **-2.40** | **0.020** | **0.098** |
| Corrected Model:  Prop. High Pos. | 0.740 | F(3,53) = 54.18 | <0.001 |  |  |  |  |
| Group |  | F(1,53) = 20.82 | <0.001 | 59.07 | 4.56 | <0.001 | 0.282 |
| Prop. High Pos. |  | F(1,53) = 4.35 | 0.042 | 10.45 | 0.43 | 0.670 | 0.003 |
| Group * Prop. High Pos. |  | F(1,53) = 2.09 | 0.154 | 47.08 | 1.45 | 0.154 | 0.038 |

Reference group = Group 1 (ASD)

**Table S3.** Pearson’s correlations between dyn-ICA and connectivity state indices in the replication sample.

|  | | **Dyn-ICA** | **p-values**  **(FDR-corrected)** |
| --- | --- | --- | --- |
| Prop. High Neg. | Pearson Correlation | .465^**^ |  |
|  | Sig. (2-tailed) | .000 | .000 |
| Prop. Moderate Neg. | Pearson Correlation | .326^*^ |  |
|  | Sig. (2-tailed) | .012 | .022 |
| Prop. Low-Uncorrelated | Pearson Correlation | -.002 |  |
|  | Sig. (2-tailed) | .988 | .988 |
| Prop. Moderate Pos. | Pearson Correlation | -.286^*^ |  |
|  | Sig. (2-tailed) | .030 | .045 |
| Prop. High Pos. | Pearson Correlation | -.358^**^ |  |
|  | Sig. (2-tailed) | .006 | .012 |
| MDT High Neg. | Pearson Correlation | .549^**^ |  |
|  | Sig. (2-tailed) | .000 | .000 |
| MDT Moderate Neg. | Pearson Correlation | .172 |  |
|  | Sig. (2-tailed) | .196 | .210 |
| MDT Low-Uncorrelated | Pearson Correlation | -.248 |  |
|  | Sig. (2-tailed) | .061 | .083 |
| MDT Moderate Pos. | Pearson Correlation | -.210 |  |
|  | Sig. (2-tailed) | .113 | .130 |
| MDT High Pos. | Pearson Correlation | -.246 |  |
|  | Sig. (2-tailed) | .062 | .077 |
| PT High Neg. | Pearson Correlation | .431^**^ |  |
|  | Sig. (2-tailed) | .001 | .003 |
| PT Moderate Neg. | Pearson Correlation | .412^**^ |  |
|  | Sig. (2-tailed) | .001 | .003 |
| PT Low-Uncorrelated | Pearson Correlation | .318^*^ |  |
|  | Sig. (2-tailed) | .015 | .025 |
| PT Moderate Pos. | Pearson Correlation | -.477^**^ |  |
|  | Sig. (2-tailed) | .000 | .000 |
| PT High Pos. | Pearson Correlation | -.422^**^ |  |
|  | Sig. (2-tailed) | .001 | .003 |

**Table S4.** Pairwise comparisons of Mean Dwell Time (MDT) in each connectivity state.

| **Group(I)** | **Group(J)** | **Mean Difference (I-J)** | **Sig.**  **(LSD adjusted)** |
| --- | --- | --- | --- |
| High negative | Moderate negative | 3.919 | .000 |
|  | Low uncorrelated | 1.706 | .003 |
|  | Moderate positive | 3.715 | .000 |
|  | High positive | -2.761 | .004 |
| Moderate negative | High negative | -3.919 | .000 |
|  | Low uncorrelated | -2.212 | .000 |
|  | Moderate positive | -.204 | .171 |
|  | High positive | -6.680 | .000 |
| Low uncorrelated | High negative | -1.706 | .003 |
|  | Moderate negative | 2.212 | .000 |
|  | Moderate positive | 2.009 | .000 |
|  | High positive | -4.467 | .000 |
| Moderate positive | High negative | -3.715 | .000 |
|  | Moderate negative | .204 | .171 |
|  | Low uncorrelated | -2.009 | .000 |
|  | High positive | -6.476 | .000 |
| High positive | High negative | 2.761 | .004 |
|  | Moderate negative | 6.680 | .000 |
|  | Low uncorrelated | 4.467 | .000 |
|  | Moderate positive | 6.476 | .000 |

**References (Supplementary Material)**

1. Alaerts, K., Woolley, D. G., Steyaert, J., Di Martino, A., Swinnen, S. P., & Wenderoth, N. (2014). Underconnectivity of the superior temporal sulcus predicts emotion recognition deficits in autism. *Social cognitive and affective neuroscience*, 9(10), 1589–1600. <https://doi.org/10.1093/scan/nst156>
2. Behzadi, Y., Restom, K., Liau, J. & Liu, T. T. (2007). A Component Based Noise Correction Method (CompCor) for BOLD and Perfusion Based fMRI. *NeuroImage*, 37(1), 90-101. <https://doi.org/10.1016/j.neuroimage.2007.04.042>
3. Díez-Cirarda, M., Strafella, A. P., Kim, J., Peña, J., Ojeda, N., Cabrera-Zubizarreta, A. & Ibarretxe-Bilbao, N. (2018). Dynamic functional connectivity in Parkinson’s disease patients with mild cognitive impairment and normal cognition. *NeuroImage: Clinical*, 17, 847-855. <https://doi.org/10.1016/j.nicl.2017.12.013>
4. Kaiser, R. H., Whitfield-Gabrieli, S., Dillon, D. G., Goer, F., Beltzer M., Minkel, J., Smoski, M., Dichter, and G., Pizzagalli, D. A. (2016). Dynamic resting-state functional connectivity in major depression. *Neuropsychopharmacology*, 41, 1822-1830. <https://doi.org/10.1038/npp.2015.352>
5. Wylie, K. P., Tregellas, J. R., Bear, J. J. Legget, K. T. (2020). Autism Spectrum Disorder Symptoms are Associated with Connectivity Between Large‑Scale Neural Networks and Brain Regions Involved in Social Processing. *Journal of Autism and Developmental Disorders*, 50(8), 2765-2778. <https://doi.org/10.1007/s10803-020-04383-w>
